# Mapping metal accumulation sites in crop fruits revealed a functional iron reservoir in the tomato seed chalaza

**DOI:** 10.1101/2023.12.12.571343

**Authors:** Kumbirai Deon Mandebere, Utku Deniz, Katarina Vogel Mikus, Seckin Eroglu

**Affiliations:** Department of Biological Sciences, Middle East Technical University, Ankara, Turkey; Jozef Stefan, Jamova cesta 39, Ljubljana, Slovenia; Biotechnical Faculty, University of Ljubljana, Jamnikarjeva ulica 101, Ljubljana, Slovenia

**Keywords:** fruit, synchrotron, X-ray, metal, iron, germination, tomato

## Abstract

Global fruit production suffers from pre- and post- harvest losses, part of which are related to metal deficiencies. Despite fruits being one of the most widely consumed plant parts, the spatial distribution of metals and their physiological significance remained largely unexplored. In this study, we investigated metal accumulation sites in fruits of 28 major crops by using X-ray and histochemical-based techniques. We found that calcium accumulated in the outermost hardened tissues, potassium in sugar accumulating fleshy tissues, and iron (Fe) in vascular tissues in a conserved manner. Vascular Fe pattern traced to the seed revealed an Fe reservoir at the fruit-seed juncture in tomato, which persisted in the seed’s chalazal region upon dispersal. To determine the physiological function of these stored reserves, we manipulated Fe bioavailability. Two opposely acting chelators, desferoxamine, an Fe immobilizer, delayed germination, while nicotianamine, a mobilizer, accelerated it in wild-type plants but not in mutants with low chalazal Fe. Additionally, external Fe supplementation also increased germination speed in a dose-dependent manner. Collectively, these findings demonstrate that fruit vasculature serves as a critical delivery system to establish seed Fe pools, which are determinants of seed germination speed. This study provides the first comprehensive atlas of metal hotspots in fleshy fruits and links anatomical distribution to a defined physiological mechanism in seed biology.

## Introduction

Fruits are essential to a nutritious human diet. To meet and sustain the dietary needs of the growing global population, fruit production must increase from one billion tons to over two billion tons per year by 2050 (Stratton et al., 2021; Willett et al., 2019). A key challenge in achieving this goal is reducing pre- and post-harvest losses, which can account for up to half of the production, making fruits the most wasted type of food (Gustavsson et al., 2011; Wang et al., 2022). Metal deficiencies are significant factors contributing to such losses, limiting the quality and quantity of fruits (Andresen et al., 2018; Kaya and Higgs, 2002; Suzuki et al., 2003). The effects of metal deficiencies can be either direct or indirect. As a direct effect, insufficient calcium levels in apples can result in bitter pit disease (Song et al., 2018; Torres et al., 2021; Van Goor, 1971). As an indirect (i.e., systemic) effect, the lack of a particular metal in sink tissues can cause metabolic remodeling, leading to lower quantity and quality of fruits (Àlvarez-Fernàndez et al., 2006; Álvarez-Fernández et al., 2003; El-Gioushy et al., 2021). Furthermore, metal deficiencies may also weaken the defense of fruits against several microbial diseases (Conway et al., 2001; Morina and Küpper, 2022).Despite the importance of metals in fruit biology, protocols for investigating their distribution in plant tissues often fail to yield the desired results.

The primary methods for exploring the spatial localization of metals in plant organs such as seeds, roots, or leaves are X-ray-based analysis and histochemical staining (Mijovilovich et al., 2020; Wu and Becker, 2012). However, these methods are often impractical for fleshy fruits with high water content (Lombi et al., 2011; Yadav et al., 2021; Zhao et al., 2014). Consequently, spatial metal analysis of fruits is currently limited to non-fleshy varieties such as grains or pods and, with only a few exceptions, does not include juicy, fleshy fruits like tomatoes or oranges (Trebolazabala et al., 2017). Mapping metals in fruits will also contribute to seed biology because the fruit is the maternal environment that loads the seed.

Fruits use complex strategies to ensure dispersed seeds germinate in favorable conditions. In a recent study, it was shown that seeds are sealed with additional hydrophobic barriers such as lignin and suberin when mother plants are subjected to high heat during the bolting stage (Hyvärinen et al., 2025). Mother plant may determine germination speed by differential loading of metals to the seeds. Plants cultivated in metal-deficient basic soils produced late-germinating seeds (Murgia and Morandini, 2017). Since Fe and copper can impact cell wall integrity through redox reactions (Chen et al., 2023) it was speculated that they may also facilitate germination by weakening the cell walls of the endosperm (Mari et al., 2020); however, such has never been directly tested. Therefore, although there are some indications, a direct link between metals and germination speed has not been established.

Here, we systematically investigated metal accumulation sites in freeze-dried fruit samples using X-ray fluorescence and histochemical staining. This approach identified metal “hot spots”, some of which are well-conserved. We further studied one of these metal hotspots to investigate its possible physiological functions.

## Material and Methods

### Research material

To examine metal localization in fruits, we selected 28 species to represent the major fruit crops consumed globally. Samples were obtained from markets and bazaars to ensure that the findings reflect metal localization patterns in agriculturally relevant, commercially available produce, rather than solely in model organisms grown under artificial laboratory conditions. Fruits were collected in Ankara, Turkey, throughout the year (Table S1). For experiments with tomato seeds, if not otherwise indicated, wild-type *Solanum lycopersicum* cv. Bonner Beste plants and its mutant *chloronerva* were used (Herbik et al., 1996).

### Tabletop XRF

To prepare the fruits for XRF analysis, they were freshly sliced both longitudinally and cross-sectionally to a thickness of 2-3 mm using a V-blade (Fig. S1). The prepared fruit slices were immediately immersed in liquid nitrogen and kept frozen at -80°C for 24 hours. Frozen samples were then placed in a vacuum freeze dryer for a main drying phase of 23 hours at -54°C and 0.024 millibars, followed by an additional hour of final drying at -60°C and 0.011 millibars. Once dried, fruit samples were stored at room temperature until the analysis.

Freeze-dried fruit samples were analyzed by a micro XRF analyzer (Horiba XGT-9000, Japan). The benchtop system was equipped with a Rh tube (50 kV, 1 mA) yielding a polychromatic excitation beam focused to 15 µm by a polycapillary lense. X-ray fluorescence was detected with a silicon drift diode (SDD) detector. The samples were mapped with 100-120 µm step size and 200 ms dwell time. The spectra in each pixel were recorded with the Horiba software, exported as .H5 files, and processed with the PyMCA 5.8.1 toolkit (Bolte et al., 2011) where full spectra deconvolution and fitting was performed. Maps were then generated in the RGB correlator of PyMCA on the basis of the intensities of particular fluorescence K lines (Solé et al., 2007).

### Synchrotron XRF

For examining metal enrichment in the vasculature of tomato fruits (*Solanum lycopersicum* cv. Moneymaker) in high resolution and with minimum treatments, fresh cut tomato pieces were embedded in the cryo-embedding matrix OCT and immediately plunged into liquid nitrogen. Fifty µm cross sections were obtained using a Leica RM2265 semi-automated rotary cryomicrotome and immediately transferred to a sample holder between two Ultralene foils. The samples were kept in liquid nitrogen during the transfer and analysis with the Scanning X-ray Microscope (SXM) at ID21 X-Ray Microscopy Beamline at the European Synchrotron Radiation Facility (Grenoble, France). The beam size was focused using Kirckpatrick Baez mirror optics to about 1×0.6 (HxV) μm^2^. Elemental localization and distribution of Fe were performed at 7.3 keV excitations above the Fe K-edge and 100 ms dwell time. The energy was selected using a double-crystal monochromator with a Si111 crystal pair. The X-ray fluorescence signal from the samples was detected using a silicium drift diode detector with an 80 mm^2^ active area from SGX Sensortech. The incoming beam intensity was monitored using a photodiode. The X-ray fluorescence data from each pixel in the maps were fitted to obtain the final elemental distribution images using the batch processing routines from PyMCA 5.9.2.

For examining metal accumulation in the chalaza of tomato seeds in high resolution, two-dimensional SXRF imaging was conducted. Dry seeds were placed on a Kapton tape. Images were collected at beamline 4-BM of the National Synchrotron Light Source-II (NSLS-II; Brookhaven National Laboratories, Upton, NY, USA). The elemental maps were collected using an incident energy of 12 keV, a 7μm step size, and a 100 ms dwell time. Raw data from the NSLS-II was analyzed using the Larch software package to produce fitted X-ray fluorescence images. Final SXRF images and intensity profile plots were generated using the Nikon NIS-elements software.

### Histochemical staining of fruits

For preparing the fruit samples for histochemical staining, freeze-dried fruit pieces that were prepared as described above for the tabletop XRF analyses were used as starting material. Colorful fruits were subjected to decolorization. Decolorization was conducted by immersing the pieces into a fixing solution (methanol: chloroform: glacial acetic acid; 6:3:1) in a glass Petri dish and shaking at 90 rpm until the natural color disappeared. The fixing solution was renewed several times during the shaking. To stop the decolorization step, the fixative was removed by washing the samples at least three times with distilled water.

For the histochemical staining of Fe, Perls staining was used as previously described (Roschzttardtz et al., 2009). Briefly, Perls stain solution was prepared fresh by mixing 4% K-ferrocyanide (K_4_Fe(Cn)_6_) and 4% HCl stock solutions in a 1:1 (v:v) proportion and vacuum infiltration for one hour at 500 millibars. Next, the staining solution was removed, and the samples were washed three times with distilled water for 1-2 min each by slight shaking. The samples were stored in distilled water at 4°C until observation with a stereo microscope.

For histochemical Fe staining on the thin cross sections of tomato fruits, samples prepared for Perls staining were first embedded in resin, cut, and then stained. Pieces were washed three times with 0.1 M phosphate buffer (pH 7.4) and dehydrated in successive baths of 50%, 70%, 90%, 95%, and 100% ethanol, butanol/ethanol 1:1 (v/v), and 100% butanol. Then, the pieces were embedded in Technovit 7100 resin (Kulzer, Tokyo, Japan) according to the manufacturer’s instructions and sliced into thin sections (10 μm). The sections were deposited on glass slides and incubated for 45 min in Perls stain solution. Perls solution was removed, slides were washed three times and directly observed with a compound microscope.

### Elemental analysis by ICP-OES

To quantify Fe accumulation in the chalazal part, elemental analysis on seed parts were conducted. To this end, mature tomato seeds (*Solanum lycopersicum* cv. Candela) were imbibed for 24 hours and dissected into three parts. Dissected samples were dried in an oven at 65°C for one week. Dried plant samples were finely ground with an electric coffee grinder. Approximately 0.3 g of the dried and ground plant samples were weighed and placed in microwave digestion tubes. On top of each sample, 2 ml of 30% H_2_O_2_ and 5 ml of HNO_3_ were added, and the digestion was conducted in a closed-vessel microwave system (MarsExpress; CEM Corp., Matthews, NC, USA). After cooling down sufficiently, the total sample volume was finalized to 20 ml by adding double-deionized water, and samples were filtered through analytical filter papers (Macherey-Nagel, Ø125 mm, blue band). Inductively coupled plasma optical emission spectrometry (ICP-OES) Agilent 5800 ICP-OES (Agilent Technologies, USA), was used to determine the concentrations of micronutrients in digested plant samples. The accuracy of the element analyses was validated using certified standard reference materials obtained from the National Institute of Standards and Technology (Gaithersburg, MD, USA).

### Ferric chelate reductase assay

Ferric chelate reductase activity of endosperm exudates was assessed according to a method previously described (Grillet et al., 2014). Mature tomato seeds (*Solanum lycopersicum* cv. Candela) were imbibed in water for 24 h at room temperature and cut longitudinally. Embryos were discarded, and five endosperms were pooled as one biological sample (n=3). Isolated endosperms were immersed in distilled water and allowed to release their exudates for 24 hours. The obtained exudate solution was then mixed (1:2 ratio) with ferric reductase activity assay solution (0.2 mM CaSO4, 5 mM MES (pH 5.5), 0.2 mM ferrozine, and 0.1 mM Fe-EDTA) on a 12-well plate in the presence or absence of ascorbate oxidase enzyme (1.5 AOX units ml^-1^). Following an incubation of 3 hours, endosperms were quickly dried on tissue paper and weighed. An 800 µl aliquot was taken from the remaining solution and the absorbance was measured at 562 nm. Experiments were repeated three times with at least three biological replicates and the result of one representative experiment is shown.

### Germination tests

For all germination experiments, plated tomato seeds were incubated for two days in a refrigerator for stratification and then placed in a growth chamber at 25 °C under continuous light. Germination was scored over time, with a seed considered germinated when the radicle protruded through the testa.

To determine how internal Fe availability impacts germination, germination tests were conducted with tomato (*chloronerva* and its wild type Bonner Beste) in the presence and absence of a strong Fe chelator. Seeds were sown on Petri dishes (30 seeds per dish, three replicates) filled with water agar (0.7% agar, buffered at pH 5.6 with MES) with or without DES (250 µM) or NA (5 µM). Results were analyzed by two-way ANOVA. Upon finding a significant difference between treatments, Tukey’s test was conducted to identify the time points treatments show significantly different results.

To determine the trends of how metal supplementation impacts germination speed, FeCl_2_, FeCl_3_, Fe-EDTA, Fe-PDMA (Prepared as previously described by Suzuki et al., 2024), MnCl_2_, and CuSO_4_ were added to water agar (0.7% agar, buffered at pH 5.6 with MES) at 100 µM final concentration of each. 50 seeds for each treatment were counted.

### Measurement of hydroxyl radical production

To quantify hydroxyl radical production in the seeds, a radiolabeled benzoyl probe, BzK_5_Me, was used as described by Fry et al. (2002) and Miller and Fry (2004). This tracer is excluded from the symplast of the cells and reacts specifically with the hydroxyl radical to form tritiated water, ^3^H_2_O. ^3^H_2_O release acts as a proxy for hydroxyl radical production. At least five seeds were imbibed in a 12-well plate in 2ml solutions on a shaker in a refrigerator for two days and one day at room temperature, in solutions containing the probe and the treatments (100 µM FeCl_2_, 5 µM NA or 100 µM Des).

Solutions were then filtered using Dowex 50 H^+^ beads to remove unreacted probe. The radioactivity of the filtrate was measured without and with evaporation following resuspension in scintillant buffer. The radioactivity of the evaporated ^3^H_2_O was calculated.

## Results

### Calcium, potassium, and iron accumulate in specific tissues of the fruit

By introducing an initial freeze-drying step, we determined metal localizations in fruit pieces obtained from various crop species using a bench-top X-ray machine. While X-ray mapping revealed metal localizations, it provided only limited resolution (Fig. S2). We examined metal distribution maps to identify conserved metal-enriched sites. We found that in all fruits, potassium accumulated in their fleshy (water-rich) parts, and calcium preferentially accumulated on the outer skin of the fruits (Fig. 1, Fig. S2).

**Figure 1:**
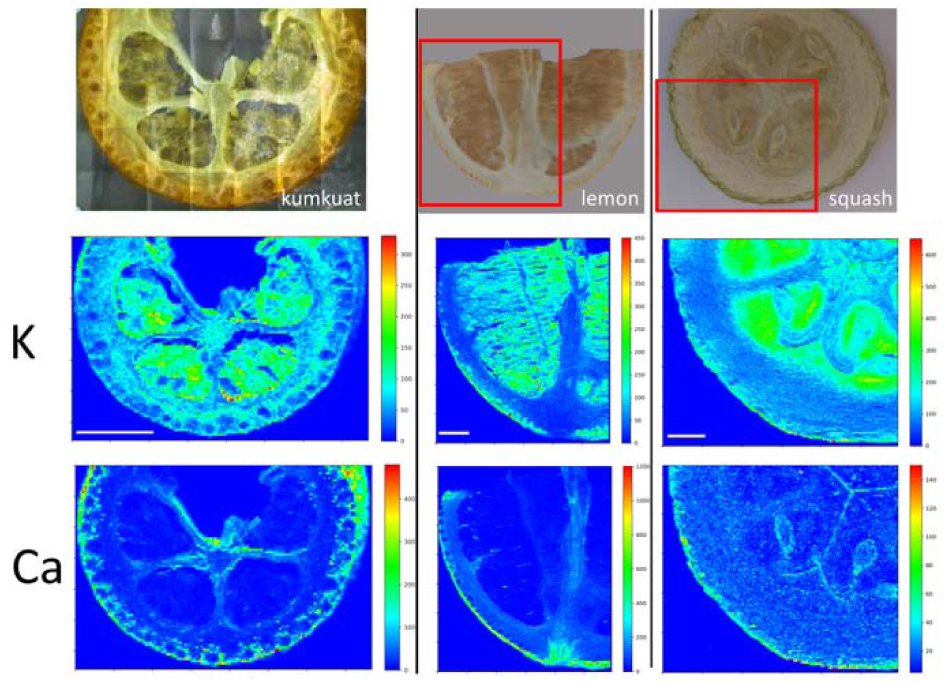
Calcium accumulates in epidermal and potassium in fleshy parenchyma tissues of fruits. Freeze-dried fruit slices were examined with a bench-top micro-XRF. Left, kumquat; middle, lemon; right, squash. Red rectangles denote the analyzed area. At least two independent pieces were observed for each fruit with comparable results. Scale bar is 5 mm. K: Potassium, Ca: Calcium.

Alternatively, histochemical stains have been used to localize metals, even in cellular compartments (Eroglu et al., 2019). To identify accumulation sites of Fe, we stained fruit slices with Prussian Blue (also known as Perls staining). Perls staining resulted in higher resolution compared to the benchtop X-ray (Fig. 2). Although Perls staining showed homogeneous coloration in some fruits (Fig. S3), it led to a more pronounced blue color throughout vascular tissues of many others (blue arrow heads in Fig. 2). This result shows that in fruits, Fe is preferentially stored in the vasculature in a conserved manner. The use of fixatives has been criticized because they may remobilize metals, leading to artifacts and misinterpretation of results (van der Ent et al., 2018). To address this constraint, we employed synchrotron X-ray mapping, which provided sufficient resolution to map vasculature in fruit tissues. Mapping of freshly cut tomato pericarp under cryo conditions revealed a hollow Fe accumulation pattern, possibly corresponding to vasculature (Fig. S4). While we were unable to identify Fe-accumulating cell types, our result suggests that the preferential Fe accumulation in the vasculature is not an artifact due to sample preparation.

**Figure 2:**
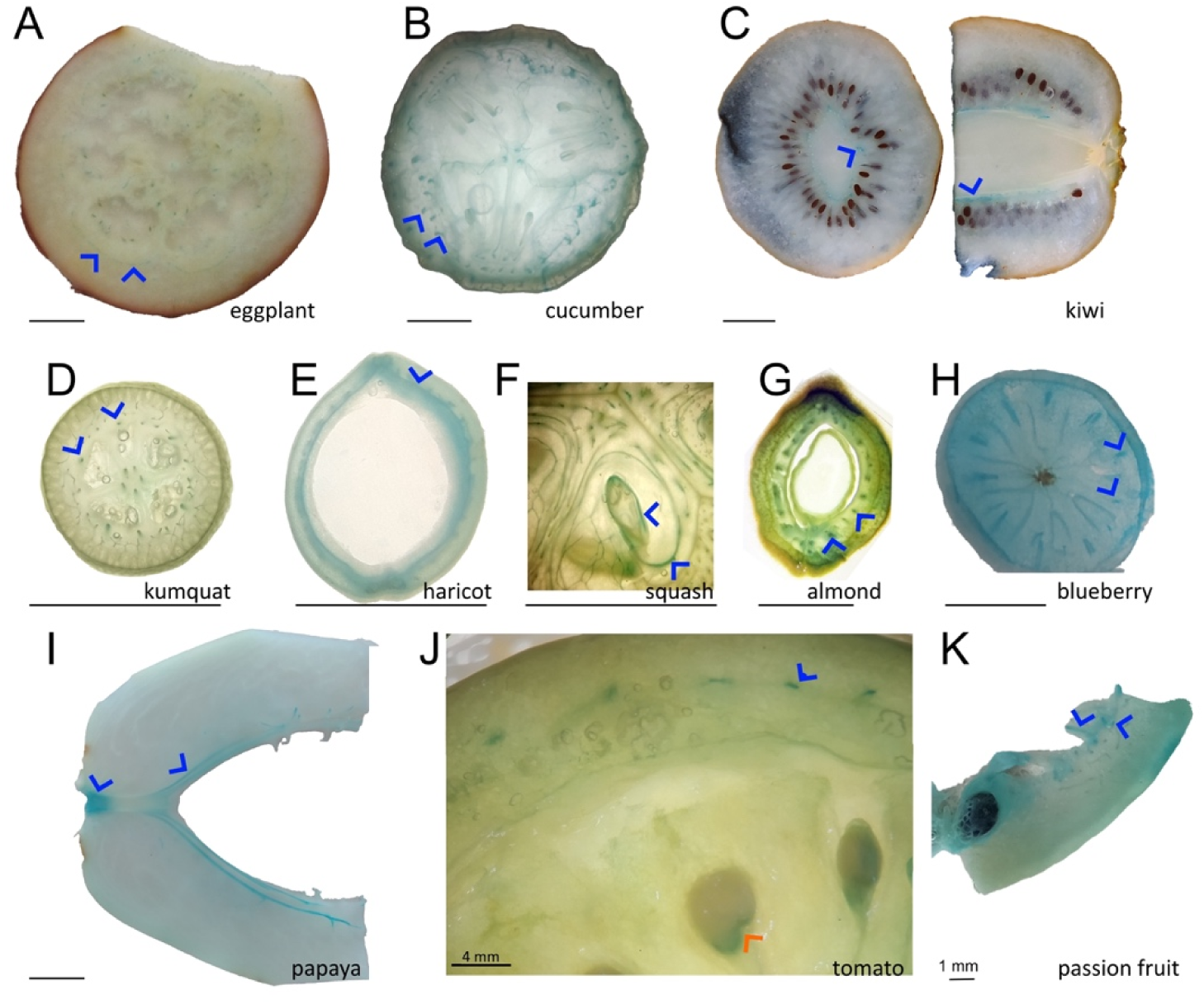
Iron accumulates in the fruit vasculature of diverse species and in the fruit-seed juncture in tomato. Cross-sectional or longitudinal cut fruit pieces were freeze-dried and decolorized with fixatives (methanol: chloroform: glacial acetic acid; 6:3:1). After fixation, the pieces were submerged in Perls staining solution for 2 to 16 hours. Blue coloration indicates Fe accumulation, blue arrowheads highlight vasculature stained blue in different fruit species. For this figure, images of fruits with noticeable Fe accumulation in the vascular bundles were chosen. Note that tomato accumulated Fe in the fruit seed juncture, shown by a red arrowhead. A-K; eggplant, cucumber, kiwi, kumquat, haricot, squash, green almond, blueberry, papaya, tomato, and passion fruit. At least five pieces of fruits were investigated, and a representative sample was photographed. Bar scale is 1 cm, unless otherwise indicated.

### Iron accumulates in the chalazal region of tomato seeds

Seeds of various species have been extensively investigated for metal distribution (Eroglu et al., 2019; Ibeas et al., 2017); however, in tomato seeds, such distribution patterns remain elusive. In the present survey, we noticed Fe accumulation in the fruit-seed juncture of tomato slices (red arrowhead in Fig. 2). We examined whether metals other than Fe are also enriched in this region by mapping seed metal distribution using synchrotron X-ray fluorescence and quantifying them with ICP-OES. Synchrotron X-Ray fluorescence showed Mn partitions towards the endosperm and Zn towards the cells that will develop into apical and root meristems (Fig. 3A). Fe accumulated both in the embryo and the endosperm. Fe accumulation in the embryo was pronounced on the prevascular strands. Fe accumulation in the endosperm was enriched in regions close to the germination site, confirming the localization obtained by Perls staining.

**Figure 3:**
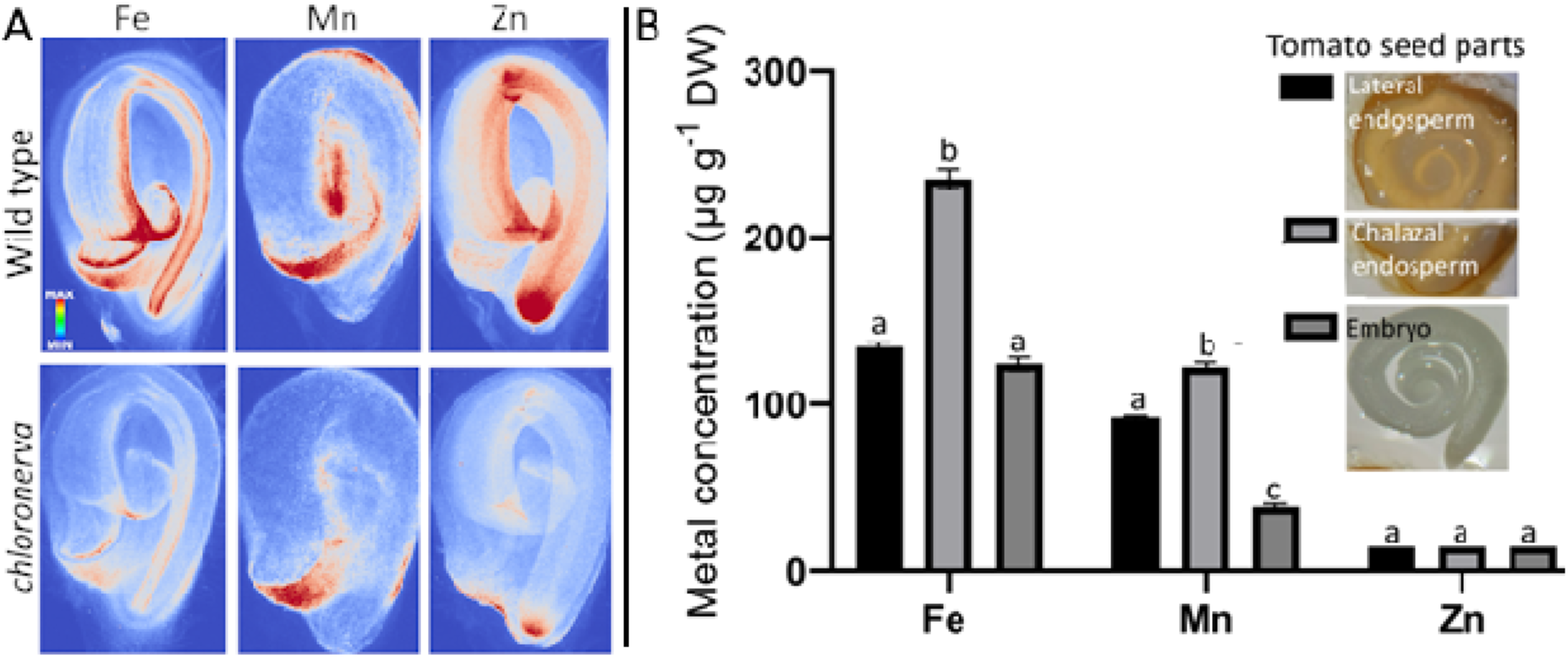
Metal loading to the fruit-seed juncture in tomato is specific to Fe and remains after seed dispersal. A, metal distribution in wild-type and mutant dry mature tomato seeds. Two seeds from each genotype were placed on Kapton tape and metal distribution was examined by synchrotron XRF. B, metal distribution in dissected seed parts. Seeds were imbibed for one day and dissected to isolate embryos and lateral or chalazal endosperm. Seed parts were dried, and metal concentrations were determined using ICP-OES. Error bars represent SEM. Different letters indicate that the means of the seed parts for that metal significantly differ from the other according to one-way ANOVA followed by a Tukey’s honestly significant difference test (*P*<0.05).

Furthermore, signal intensity in the chalaza was comparable to that of the embryo. A tomato mutant, *chloronerva,* was reported to be devoid of a Fe chelator, NA, and to contain less Fe (Herbik et al., 1996). In addition to wild-type seeds, we examined *chloronerva* seeds to assess the impact of nicotianamine on seed Fe distribution. As expected, *chloronerva* showed lower signals for all Fe, Mn, and Zn, indicating lower levels of these metals in the mutant (Fig. 3A, bottom panel). However, except for predominant Mn accumulation in the chalaza, *chloronerva* showed the same metal distribution pattern as the wild-type tomato. We also examined spatial metal distribution in the wild type seeds by analyzing metal concentrations in cut seed parts. This analysis showed the chalazal endosperm (more accurately, chalazal and micropylar endosperm, including the associated seed coat) contained approximately twice as much Fe compared to the remainder of the seed in agreement with the X-Ray fluorescent measurements and Perls staining. Overall, the mapping of metals in tomato seeds showed that the chalaza is a primary Fe-enrichment site, and *chloronerva* can serve as a tool to investigate the physiological function of this store.

In a germinating seed, endosperm-accumulated nutrients can be remobilized to support the embryo as it transforms into a seedling (Uraguchi and Fujiwara, 2011). To validate a possible role for Fe in feeding the germinating embryo, we examined the fate of the Fe deposits after germination. In seeds of one-week-old seedlings, seed endosperm tissues were degraded, reminiscent of nutrient remobilization from endosperm into the embryo (Fig. S5). However, Perls staining showed that these endosperm remnants preserved the preferential Fe accumulation in the chalaza (Fig. S5, denoted “After”) similar to the ungerminated control (Fig. S5, denoted “Before”). Metal analysis showed that the Fe concentration doubled. Since the cultivation media lacked nutrients, the observed increase in Fe concentrations was primarily due to mass loss from endosperm degradation. These results negate the hypothesis that Fe in the chalaza nourishes the germinating embryo.

### Iron is involved in germination

Next, we assessed whether Fe could promote germination by comparing the effects of Fe chelators on seeds with contrasting Fe concentrations. The Fe chelator DES has been used to counteract Fe poisoning in humans (Rafati Rahimzadeh et al., 2023) and to render Fe stores in the plant tissues unavailable for biological processes (Aznar et al., 2014). Targeting Fe stores in seeds with DES led to a delay of approximately 6 hours (Fig. 4A). Following the same logic, we also added NA to the medium. As previously mentioned, plants use NA to mobilize metals in the long distant transport (Takahashi et al., 2003). Although the effect was less striking than that of DES, NA increased germination speed(Fig. 4A). However, the germination of the *chloronerva* mutant, which contains low concentrations of Fe, was not affected by either DES or NA (Fig. 4B), suggesting that seed Fe reserve acts as a determinant for germination speed. After investigating the effect of Fe mobilization on germination speed, we examined whether Fe application can promote germination similar to NA. We added various forms of Fe, both chelated and unchelated, to the medium. Results showed that independent of the form, Fe increased germination speed (Fig. 4C). To determine whether the early germination phenotype is specific to Fe, we tested various other micronutrients. None of them caused an apparent increase in the germination speed (Fig. 4D). Interestingly, increasing Fe concentrations, even to physiologically toxic levels (200 and 500 µM), tended to increase the germination speed further (Fig. 4E).

**Figure 4:**
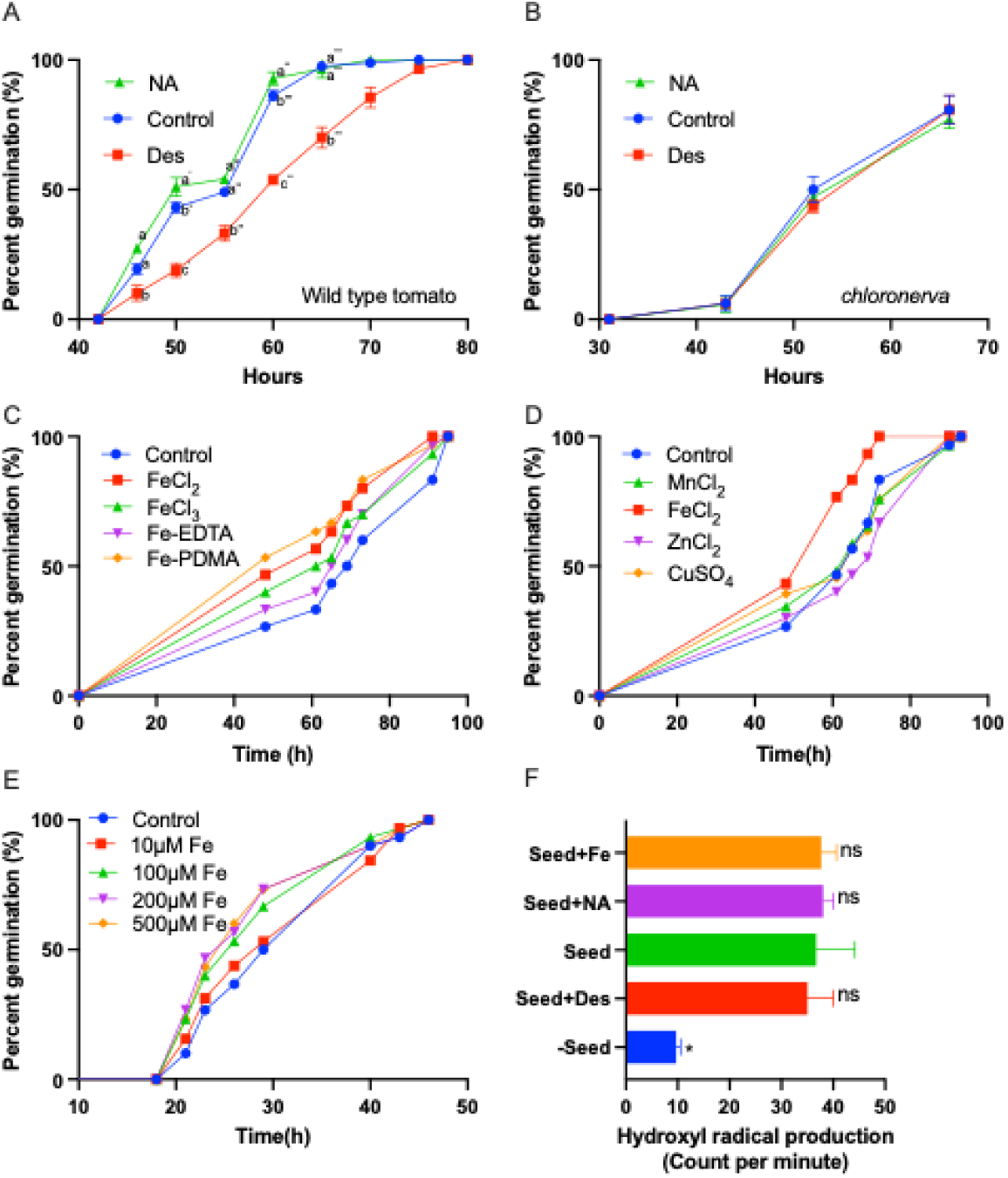
Iron facilitates germination of tomato seeds without inducing hydroxyl radicals. A, impact of chelators desferoxamine (Des)(100 µM) and nicotianamine (NA)(5 µM) on germination speed of wild type seeds. B, impact of the same chelators on germination speed of *chloronerva* mutants that contain low chalazal Fe. For A and B, Each treatment consisted of three plates, each containing at least thirty seeds (n=3). Error bars represent SEM. Different letters at the same time point indicate a significant difference determined by two-way ANOVA, followed by Tukey’s test for individual time points (*P*<0.05). The experiment was repeated four times with comparable results. C, impact of Fe form on germination speed. Final concentration of each Fe form was set to 100 µM. D, impact of several micronutrients on germination speed. Final concentration of each micronutrient was set to 100 µM. E, impact of increasing Fe-EDTA concentration on germination speed. For C-E, fifty seeds for each treatment were assessed. F, effect of Fe availability on hydroxyl radical production. Five seeds were pooled as a biological replicate (n=3). Seeds were incubated in water containing a radioactive probe, BzK_5_Me, which is reactive to hydroxyl radicals in the presence of Fe (250 µM FeCl_2_) or Fe chelators (5 µM Na or 100 µM Des). ^3^H_2_O release was considered as a proxy for hydroxyl radical production. Error bars represent SEM. The asterisk indicates that the corresponding mean of the treatment or negative control is significantly different from the mean of the control (Seeds imbibed without chelators or Fe) according to Student’s *t*-test (*P*<0.05).

Germination speed is determined by the amount of hydroxyl radicals produced in the endosperm (Muller et al., 2009; Schweikert et al., 2002). To investigate the mechanism of Fe-triggered germination, we hypothesized that Fe aids germination by weakening the endosperm through the production of hydroxyl radicals. A common pathway in nature for hydroxyl radical production is through the Fenton chain reaction, which requires Fe, hydrogen peroxide, and ascorbate (Burkitt and Gilbert, 1990a). Since, hydrogen peroxide production in tomato seeds during imbibition was shown before (Morohashi, 2002) and we found that tomato endosperm also produces ascorbate during imbibition (Fig. S6); we predicted that hydroxyl radical production to increase in the presence of Fe. Comparison of hydroxyl radical production showed that seeds produce hydroxyl radicals during imbibition, but this production was independent of Fe availability (Fig. 4F).

## Discussion

X-ray and histochemistry-based methods highlighted numerous metal hotspots in various plant organs (Eroglu et al., 2019, 2016; Kim et al., 2006). In this study, we examined metal localizations in fruits. Our results show that in fruits, metals including calcium, potassium, and Fe preferentially accumulate in certain tissues. We further focused on one of the Fe accumulation sites, in the fruit-seed juncture, and found that this Fe store mobilizes to facilitate seed germination upon seed dispersal.

### Histochemical staining of freeze-dried and decolorized fruits provides tissue-level information

The histochemical approach to mature fleshy fruits often faces two main obstacles: 1) too much water, limiting the penetration and activity of chemicals, and 2) too many pigments, limiting the visual inspection of the stain. Industrially, fruits have been extensively dried for various purposes. Among drying technologies, vacuum freeze drying appears to be the golden standard for human consumption (Harper and Tappel, 1957). We observed that once freed from water, the fruit slices behaved like regular tissue. After vacuum freeze drying, fixatives could decolorize the fruit. This may represent an advance over the techniques described in the literature. The most common way to circumvent the natural bright colors of fruits like tomato, has been to analyze the sample in earlier developmental stages (Saed Taha et al., 2012; Wang et al., 2014). We propose here that freeze drying, coupled with histochemistry, provides an efficient and quick method to reveal immobile metal hotspots in fruits (Fig. 2 and Fig.S3). Any finding can then be later confirmed with more advanced methods that offer higher resolution and quantification and avoid possible metal re-localization (Fig. S4).

### Iron follows the vasculature of fruits and seeds

Fe preferentially accumulates in the vascular tissues of several plant parts. Leaves (Xu et al., 2022), embryos inside the seeds (Kim et al., 2006 and Fig. S7), and fruits (this study) accumulate Fe in and around their provascular or vascular strands. In mature fruits, vascular Fe may represent the most prominent Fe accumulation site (Fig. 3).

However, this accumulation can appear more pronounced than it is, due to clearing off weakly bound Fe in the rest of the fruit during fixation and decolorization. The physiological function of Fe in the fruit vasculature is unclear. Fe deficiency in fruit trees has not been linked with a disease that is specific to fruit vasculature (Àlvarez-Fernàndez et al., 2006). Alternatively, an immobile Fe reservoir may be a “byproduct” of extensive callose production. Callose is required to close the xylem connection in fruits (Knipfer et al., 2015) and may cause the oxidation of a large amount of Fe for its production (Müller et al., 2015). Another possibility is that the main vasculatures of fruits serve as a sink to load Fe into the seeds. In this model, the vasculature may represent a reservoir of excess Fe left over from seed Fe filling, as this process is highly regulated to prevent Fe-mediated oxidative damage in seeds (Sun et al., 2021). The remaining Fe can then be bound to cell walls that can quickly absorb Fe(III) (Xu et al., 2022). The current study identified the vasculature as the main Fe-accumulating tissue in fruits; however, further investigations to identify the cells within the vasculature that accumulate Fe and into the underlying mechanisms of this process are required to substantiate this supposition.

### Iron in the seed chalaza facilitates germination

A fraction of Fe-loaded vasculature retains in the seeds upon dispersal from the fruit (Fig. 3, S5, S7 and Liu et al., 2023). This Fe reservoir facilitates germination (Fig. 4). Germination requires softening of the endosperm to allow radical protrusion, a major process that includes cell wall-cutting hydroxyl radicals (Grillet et al., 2013; Muller et al., 2009). A well-known source of hydroxyl radicals derives from the reaction of Fe with H_2_O_2_, referred to as Fenton reaction (Burkitt and Gilbert, 1990b). Therefore, Fe deposits in the chalaza may facilitate germination by generating cell wall-attacking hydroxyl radicals. This hypothesis is ruled out, as we failed to link hydroxyl radical production to Fe availability (Fig. 4F). To elucidate the mechanism, future research should integrate reverse genetic screens (especially under high Fe supplementation) to identify genes involved in Fe-triggered germination.

### Micro-XRF mapping is applicable to investigate nutrient deficiency-related fruit diseases

Micro-XRF mapping has been successfully used to examine metal distributions in various plant organs, including leaves, roots, stems, and seeds (Singh et al., 2022). We detected calcium to be preferentially accumulating in the outer part of fruits, especially in the citrus genus (Fig. 1, S2). This is in agreement with previous reports where the highest calcium levels were generally observed in the peel of the fruits (Saure, 2005).

Higher calcium in the outer parts of the fruits is probably caused by cell wall thickening. Calcium is a major component of the cell wall (Thor, 2019), and outer parts of the fruits usually possess thicker cell walls to maintain integrity (Forlani et al., 2019; Hallett and Sutherland, 2005; Jeffery et al., 2012). When fruits contain insufficient calcium they may be more prone to infections, as calcium deficiency has been reported to cause several diseases, including bitter pit in apples and blossom-end rot in tomatoes, peppers, and watermelons (de Freitas and Mitcham, 2012). Potassium preferentially accumulates in the fleshy parts of the fruits in a conserved manner (Fig. 1 and Ahmed et al., 2024). This is not surprising since potassium is involved in osmoregulation and sugar accumulation, whereas the fleshy parts of the fruits often accumulate water and sugar (Kumar et al., 2006).

Imaging element distributions of fruits is associated with certain limitations. Polychromatic excitation with an X-ray tube (Rh in our case) gave a high background due to scattering in the energy range of middle z-elements, decreasing the detection limit for trace elements like Fe, Cu and Zn. We found that in fruits, Fe was under the detection limit for imaging despite the presence of an Fe peak (Fig. S8). Nevertheless, our data showed that micro-XRF can be very useful to image distribution of macro-elements like potassium and calcium which can aid in investigating whether specific disease symptoms overlap with perturbations in metal accumulating sites.

## Conclusions

Increasing fruit production should target decreasing pre- and post-harvest losses. To reduce such losses, a better understanding of fruit biology is required. This study extends the basic techniques to localize metals to mature fleshy fruits, contributing to preventing yield loss in fruit production and storage in the long run. Fruits preferentially accumulate Fe in their vascular, potassium in their fleshy and calcium in their epidermal tissues. While tomato seeds preferentially accumulate Fe in the fruit-seed juncture, seed Fe determines germination speed.

## Author contributions

S.E. designed the study. U.D. and D.M. prepared the fruits and conducted the Perls staining. D.M. and S.E. conducted the seed-related experiments. K.V.M conducted benchtop X-Ray analyses. All authors discussed the data. S.E. wrote the manuscript with contributions from the other authors. All authors read and approved the final version of the manuscript.

## Acknowledgments

We thank Mary Lou Guerinot and Ryan Tappero for synchrotron imaging at X-ray Fluorescence Microprobe (XFM) beamline 4-BM of the National Synchrotron Light Source II, a US Department of Energy (DOE) Office of Science User Facility operated for the DOE Office of Science by Brookhaven National Laboratory under Contract DE-SC0012704. We acknowledge the European Synchrotron Radiation Facility (ESRF) for provision of synchrotron radiation facilities under proposal number LS-3343 for beamline, and we would like to thank Hiram Castillo for assistance and support in using beamline ID21. We also acknowledge the SIMBION (Eurobioimaging) network for benchtop micro-XRF analyses and ARIS for program financing (P1-0212, Plant Biology). We further thank Petra Bauer for *chloronerva* mutants, Abdulsamet Sakalar for helping with the histochemical staining of fruits, Catherine Curie for help with the germination experiments, Stephen Fry for hydroxyl radical measurements and Ahmet Bakirbas for critical reading of the manuscript. This study was supported by the Scientific and Technological Research Council of Turkiye (TUBITAK) under Grant Number 122Z109; European Union under COST action CA19116, Trace metal metabolism in plants (PLANTMETALS), EMBO scientific exchange grant (10056) and TENMAK, Ankara, Turkey.

