## supplementary figure for "Mapping metal accumulation sites in crop fruits revealed a functional iron reservoir in the tomato seed chalaza"

**Supplementary Figures**

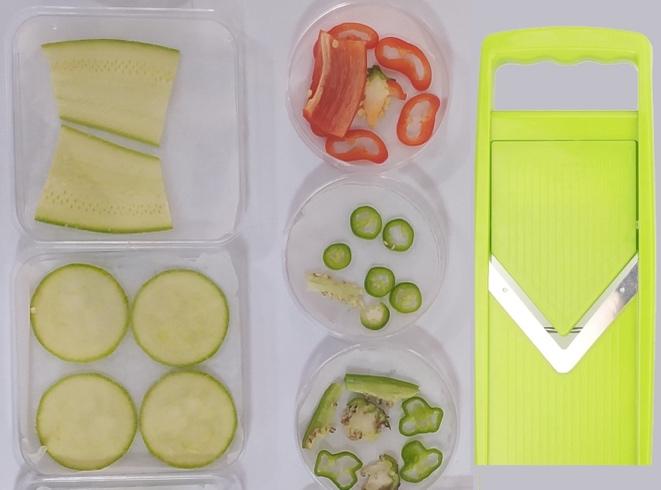

**Figure S1: V-blade cutter allows for obtaining slices of similar size from the fruits.**

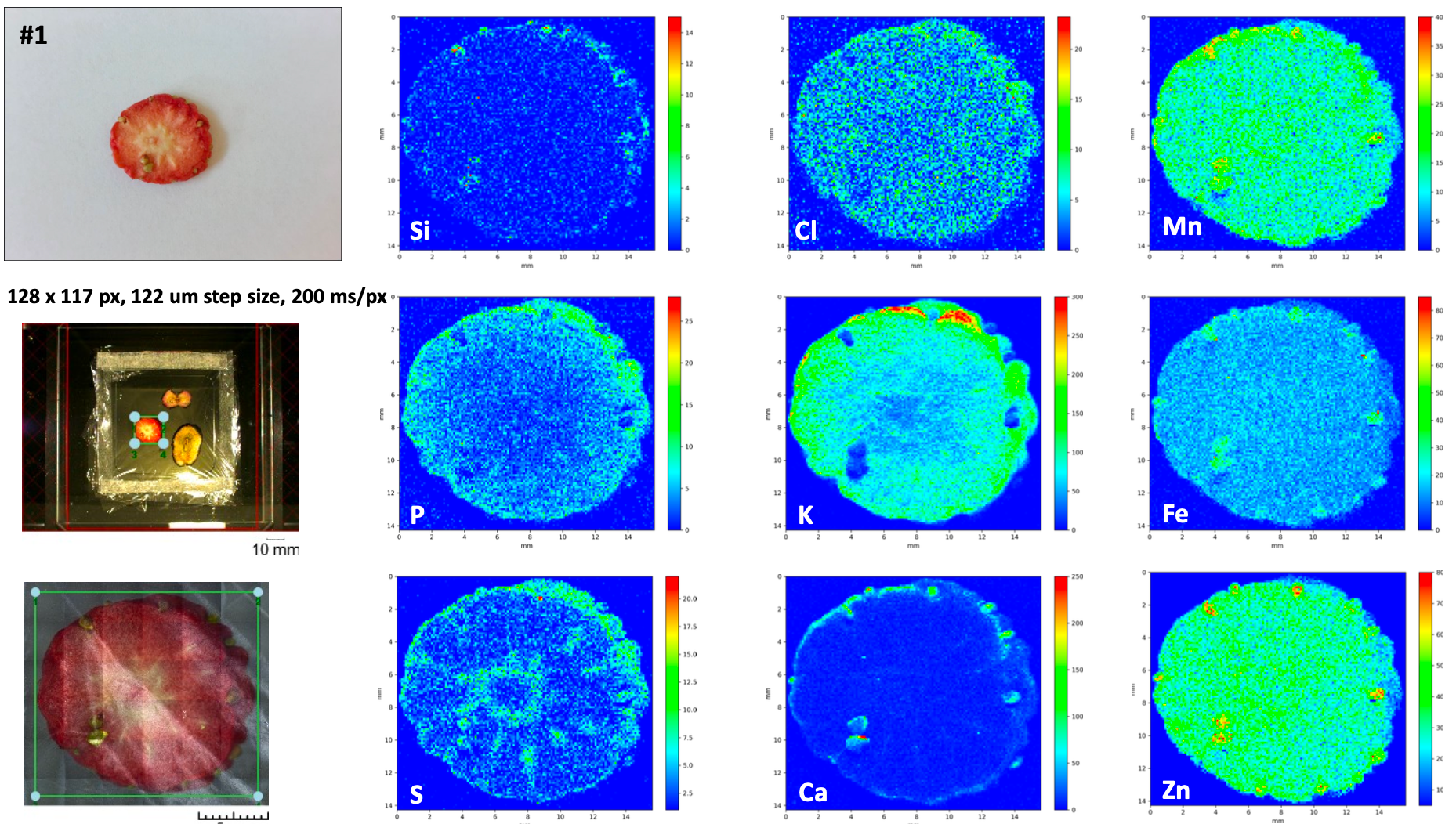

Strawberry

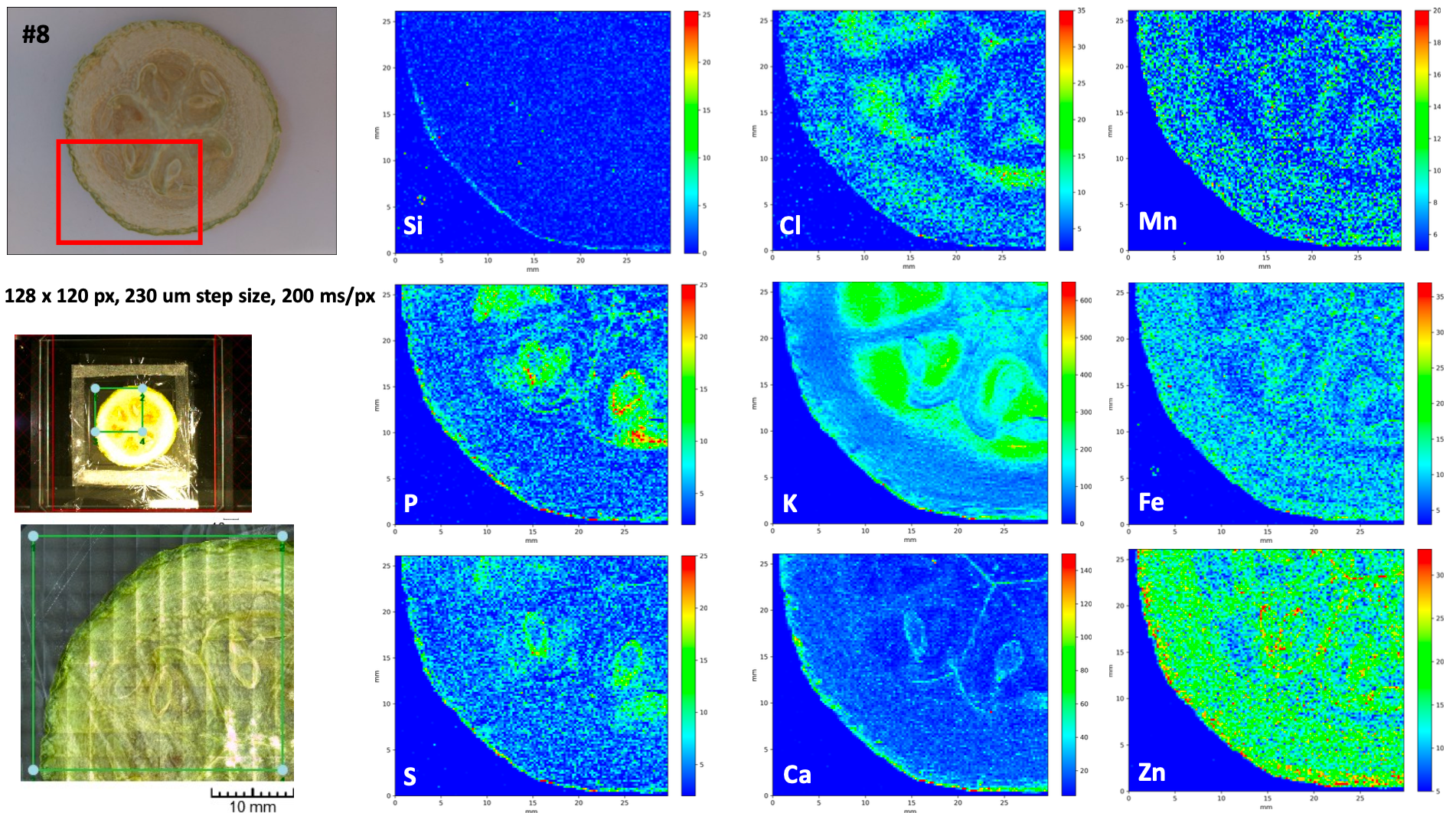

Squash

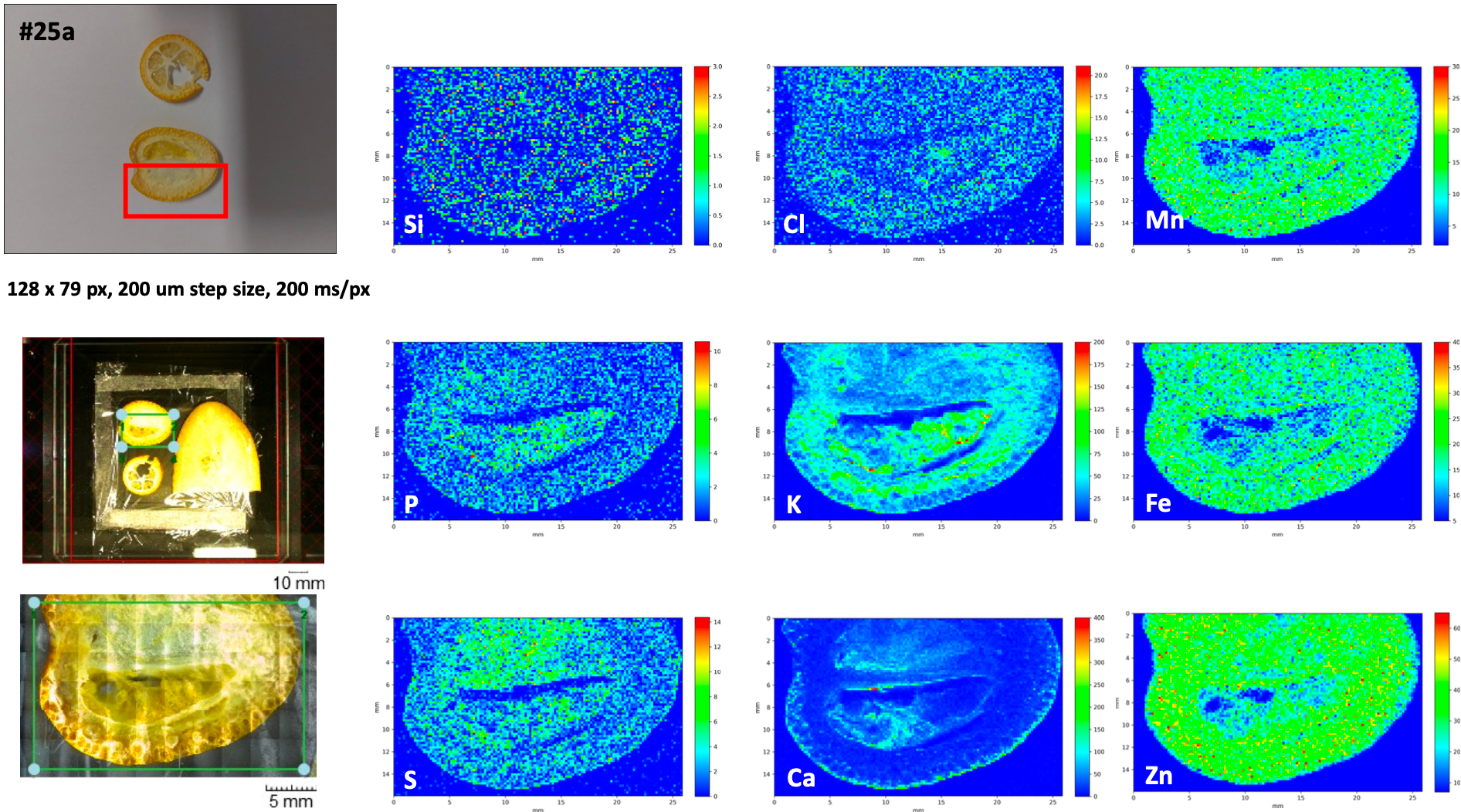

Kumkuat

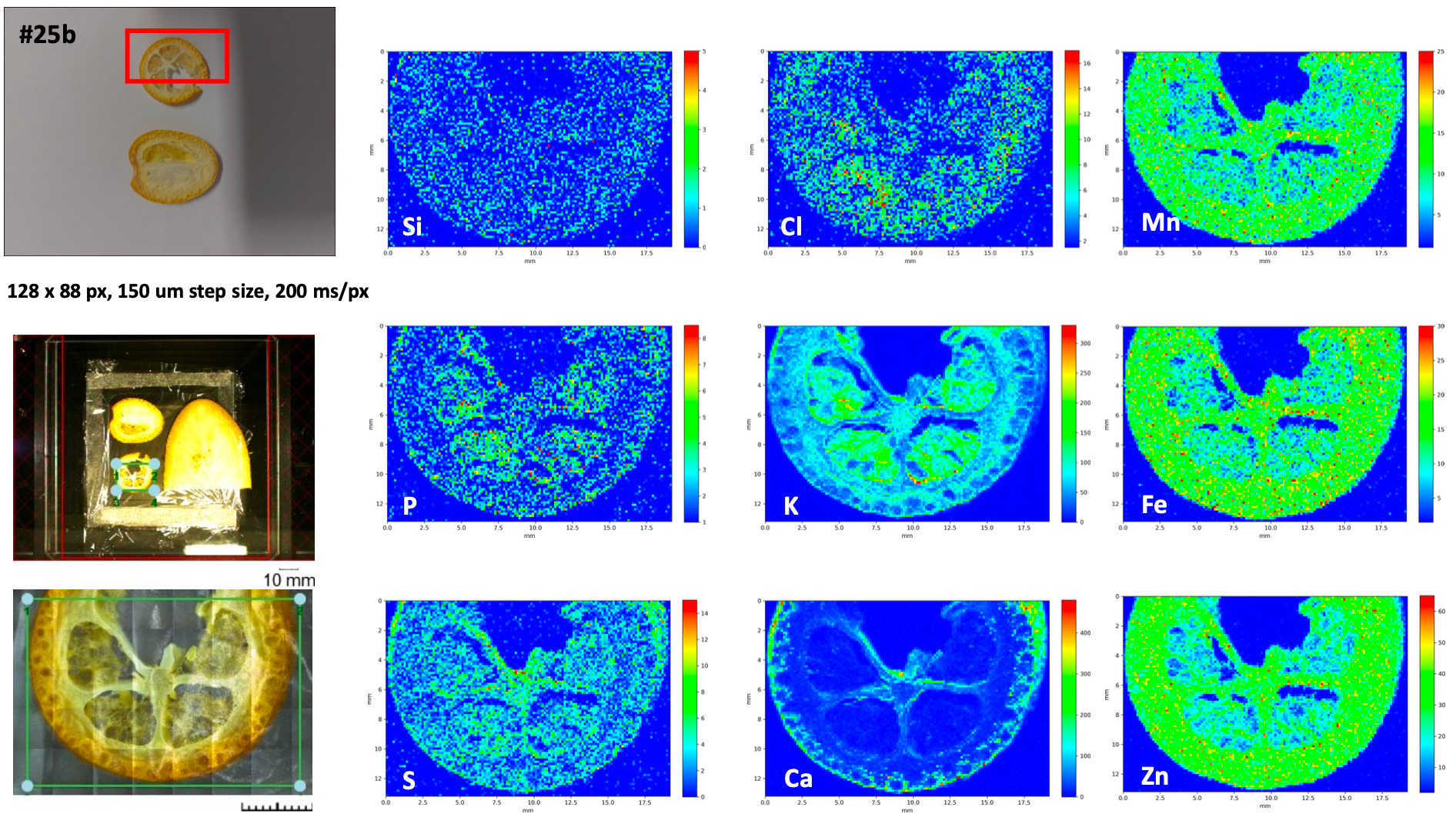

Kumkuat

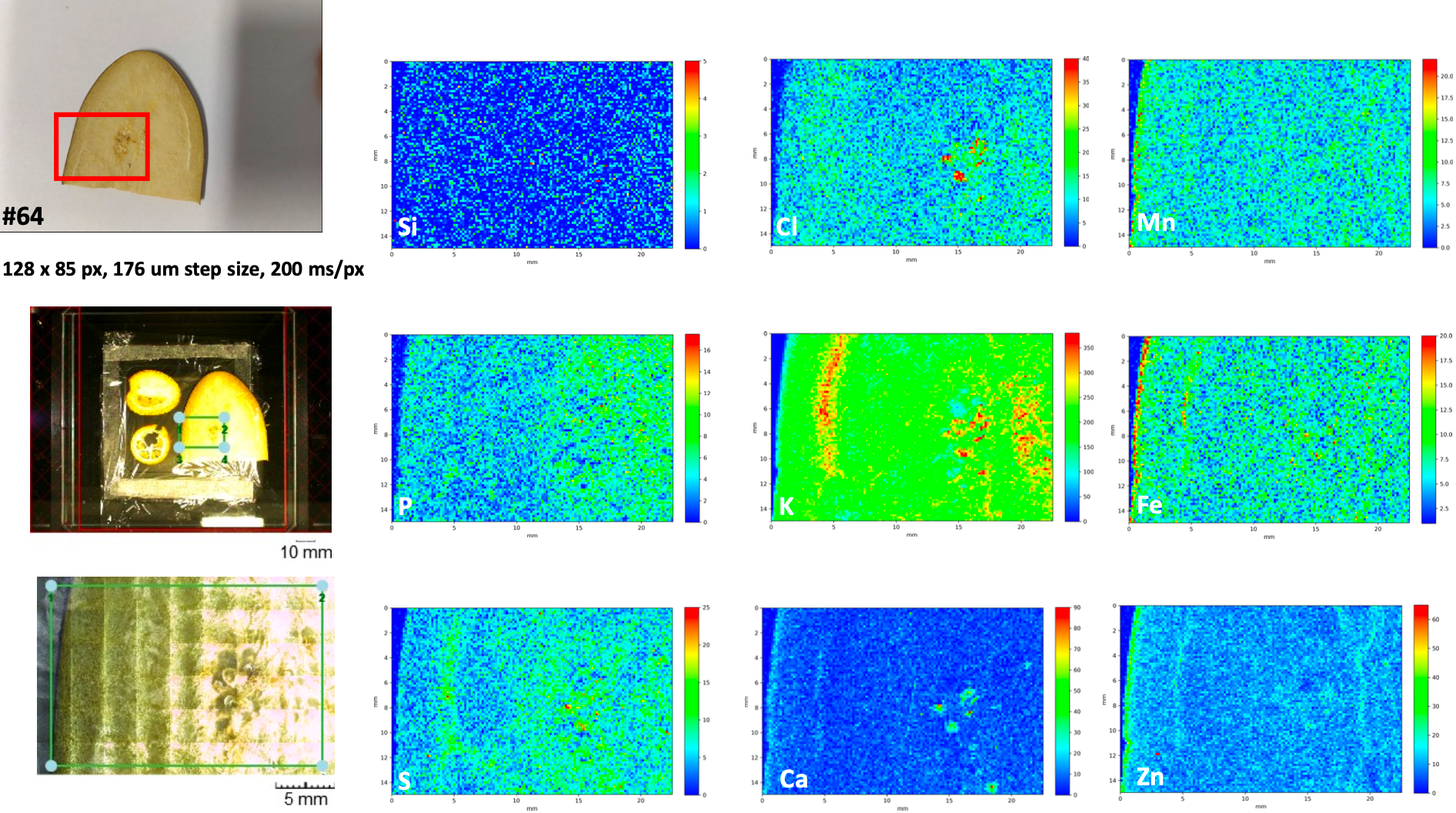

Aubergine

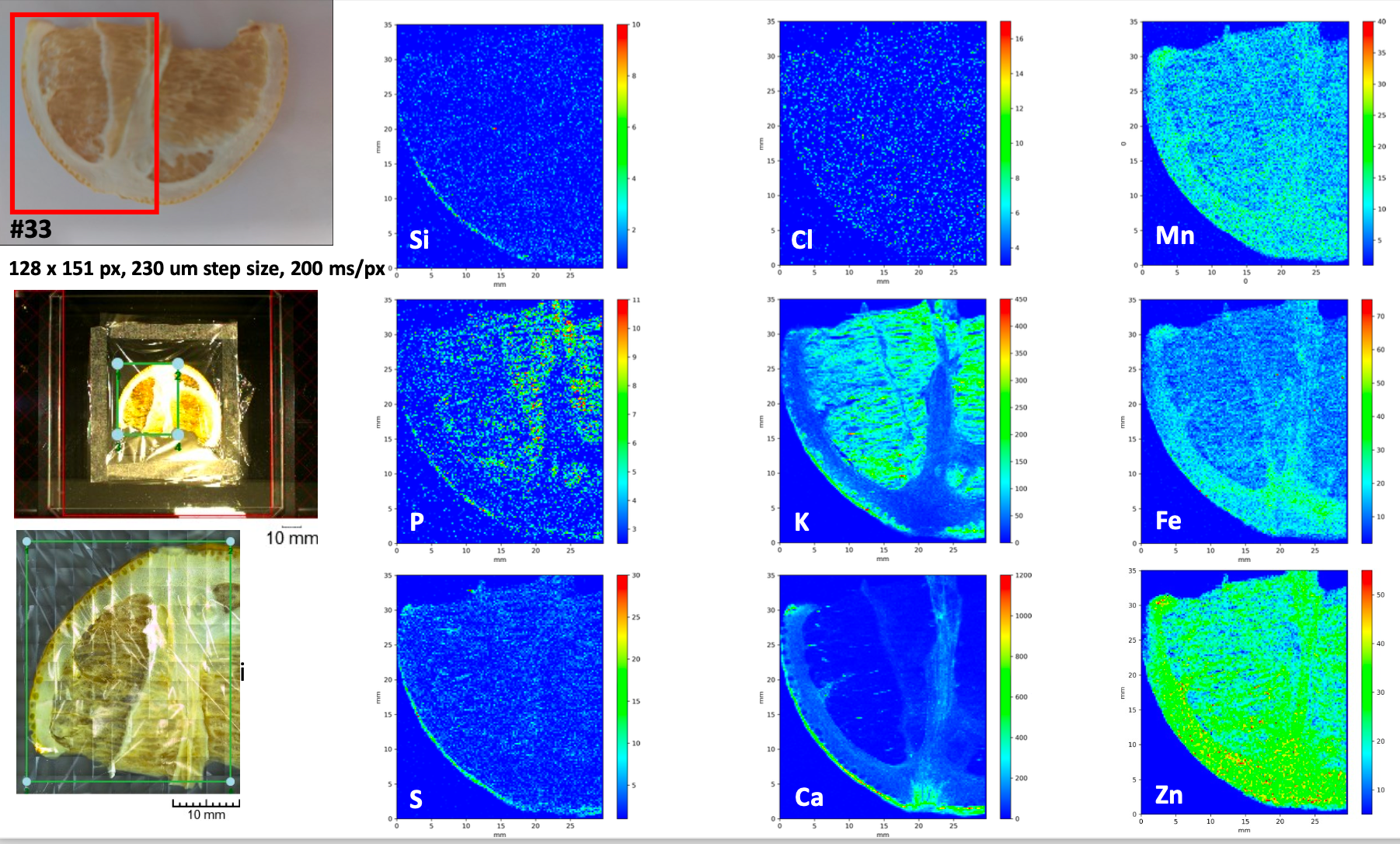

Lemon

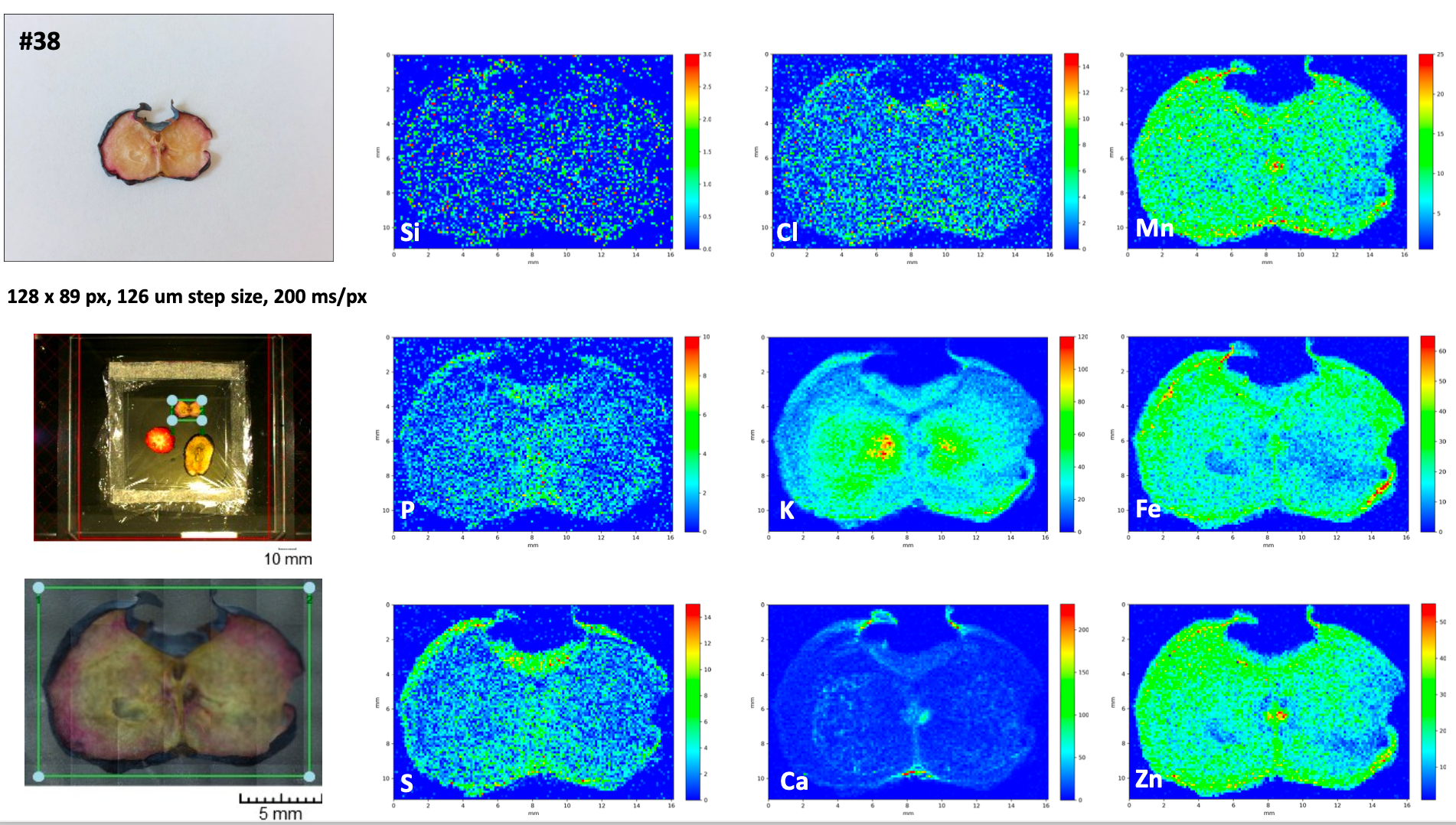

Blueberry

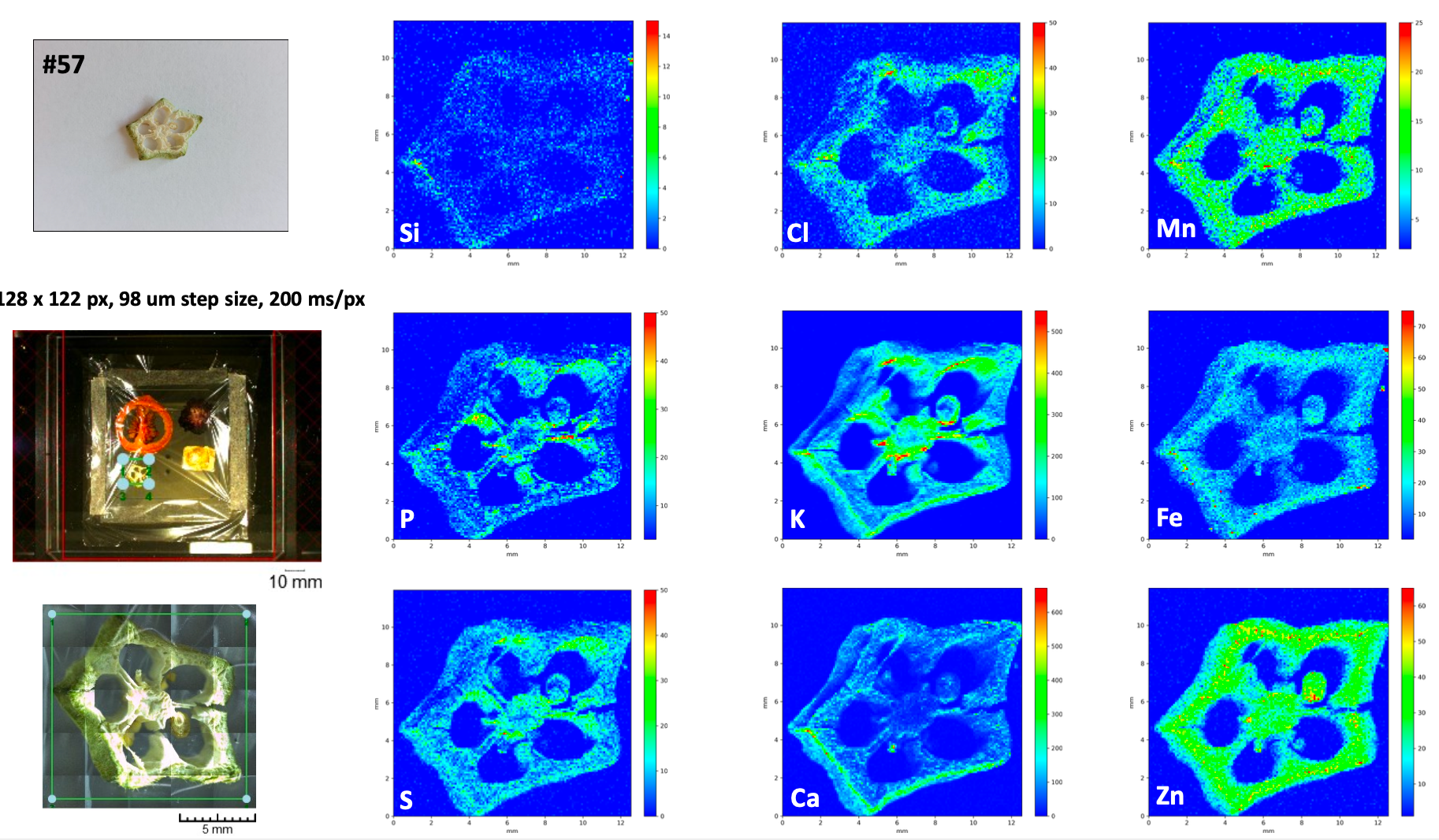

Okra

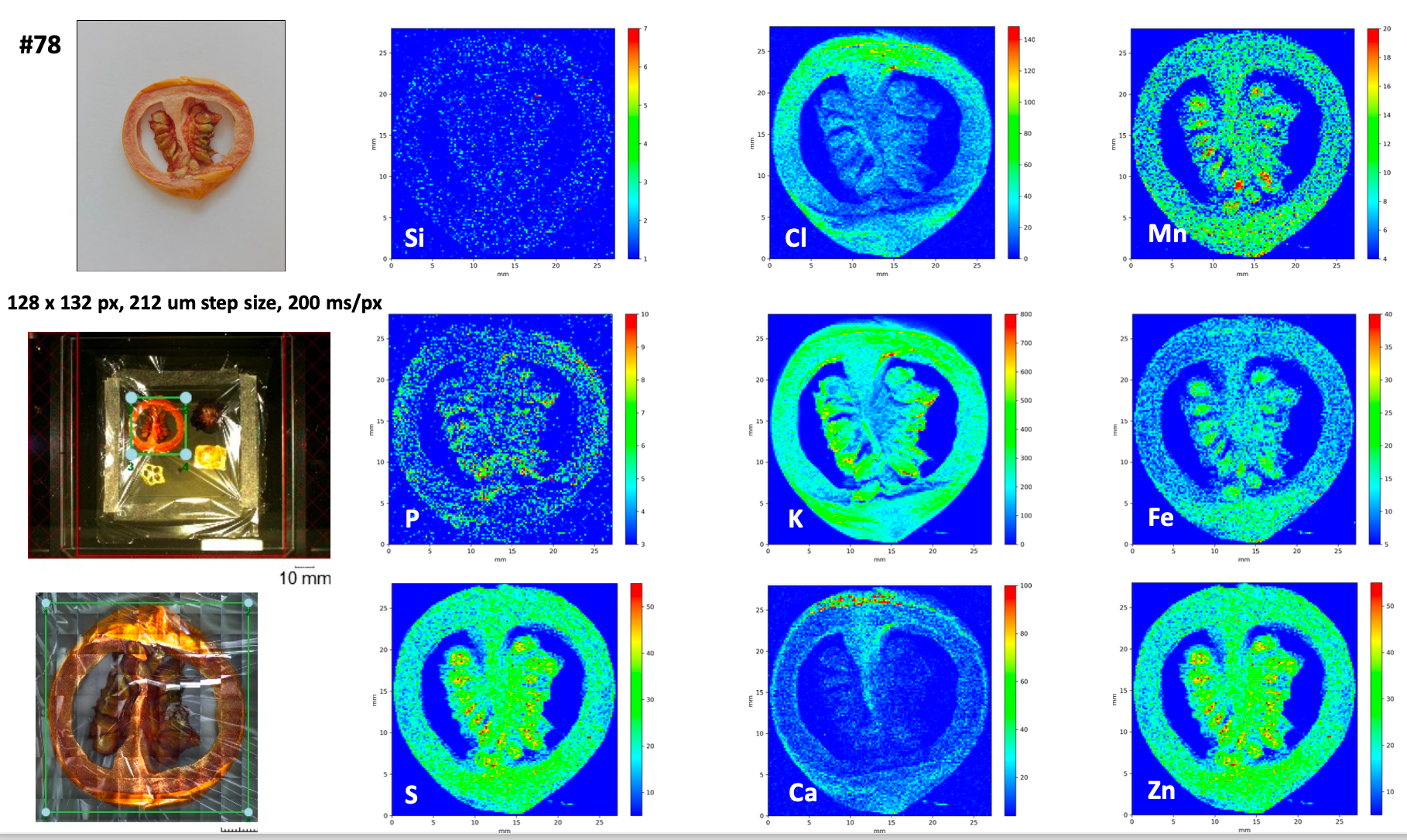

Tomato

**Figure S2: micro-XRF maps of various fruits.** From top to the bottom: strawberry, squash, kumkuat, aubergine, lemon, blueberry, okra, tomato.

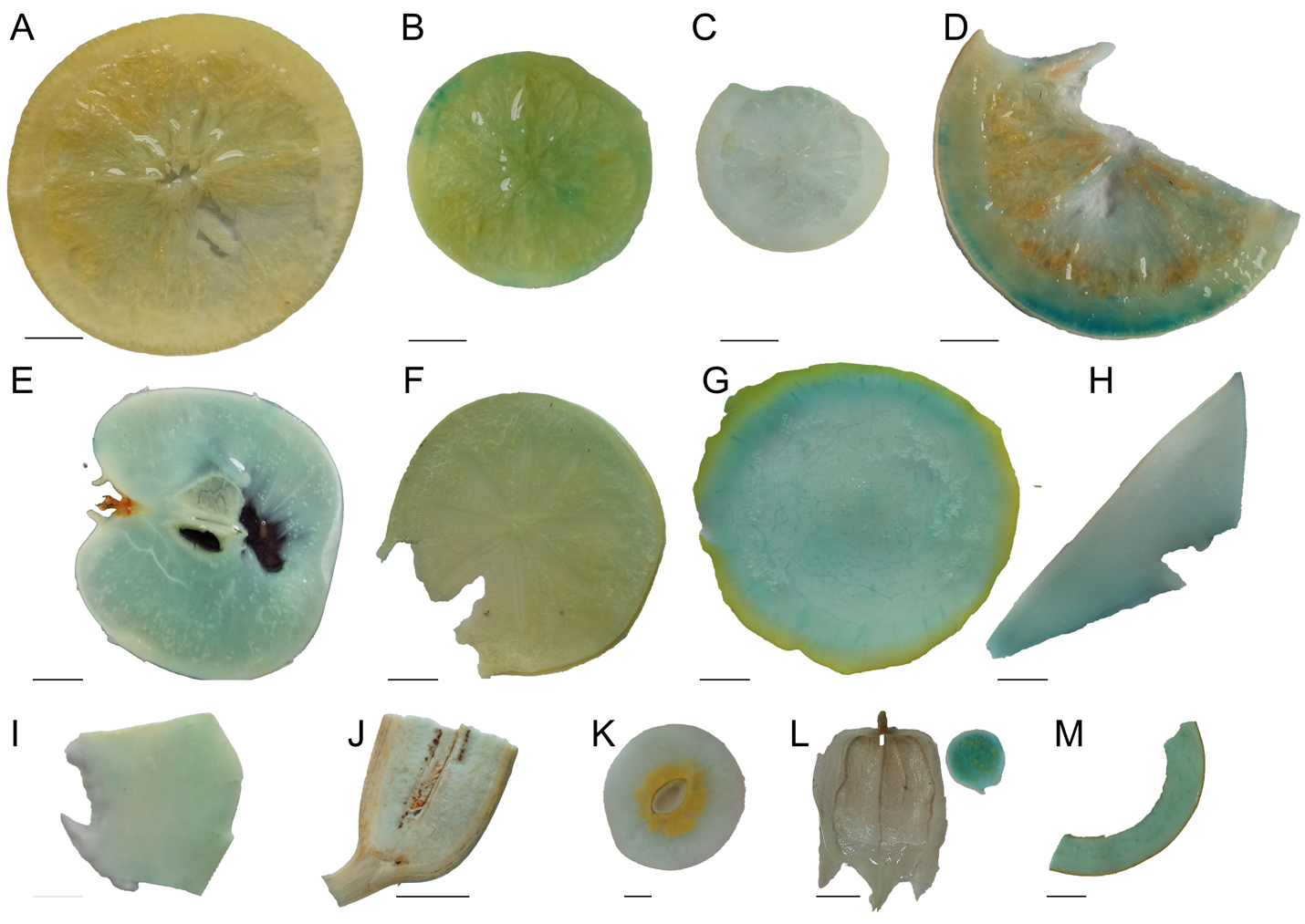

Pumpkin	Plantain	Nectarin	Cape gooseberry	Avacado

Orange	Mandarin	Lemon	Grapefruit

Apple	Persimmon	Honey melon	Mango

**Figure S3: The newly developed protocol revealed homogeneous Fe accumulation in various fruits.** Fruits were either cut cross-sectional or longitudinal with a V- blade (For the V-blade, refer to Fig. S1). These pieces were then freeze-dried and decolorized using fixatives (methanol: chloroform: glacial acetic acid; 6:3:1). After fixation, the samples were submerged in Perls staining solution between 2-16 hours. A to M; orange, mandarin, lemon, grapefruit, apple, persimmon, honey melon, mango, pumpkin, plantain, nectarin, cape gooseberry, avacado. Note that despite the differences in fruit types, the optimized protocol efficiently preserved the structure and decolorized the tissues. For the complete list of successfully stained fruits, refer to Supplementary Table 1. Blue coloration shows Fe. Bar 1 cm unless otherwise indicated.

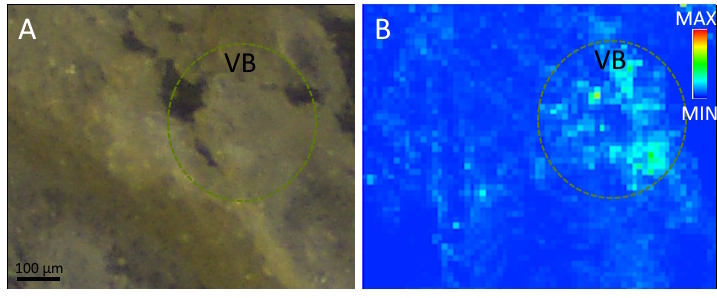

**Figure S4:** **The preferential accumulation of Fe in the vascular bundle and is not attributed to artifacts in sample preparation.** A, B; cross sections of fresh tomato fruits. Cross sections (45 µM) were prepared from outer pericarp of tomato fruits. Fresh cut tomato fruit pieces were embedded in OCT through cryosectioning. A, light microscopy image of the cross-section. B, SXRF image of the same section illustrating Fe localization. Two independent sections were generated and analyzed with comparable results. A, Light microscopy image. B, SXRF image for Fe localization. VB: vascular bundle

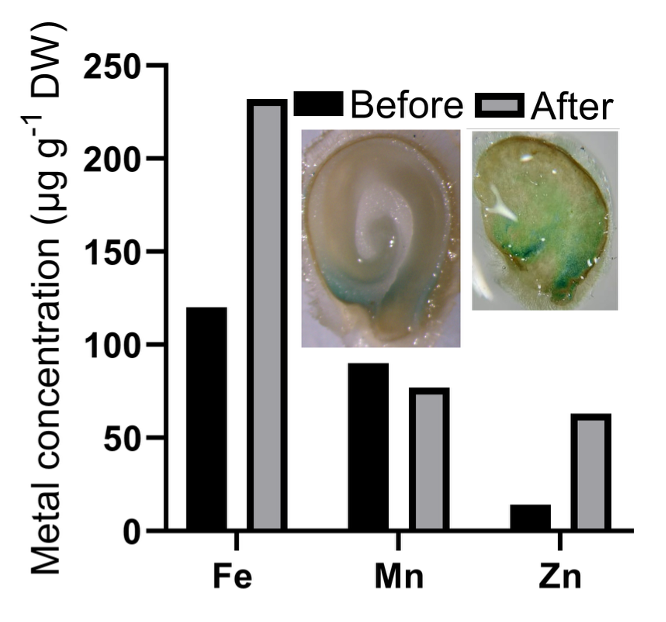

**Figure S5: The Fe hotspot in the chalazal endosperm is retained after germination.** Endosperms were isolated either from ungerminated seeds (Before, black bars) or collected from one-week-old empty seeds (leftover of established seedlings, After, grey bars). Part of the seeds were stained using Perls to disclose the Fe localization as illustrated in the pictures above the bars. The blue coloration indicates Fe accumulation. The remaining collected material (whole endosperms+seed coats of ungerminated or germinated seeds) was analyzed for metal concentrations using ICP-OES.

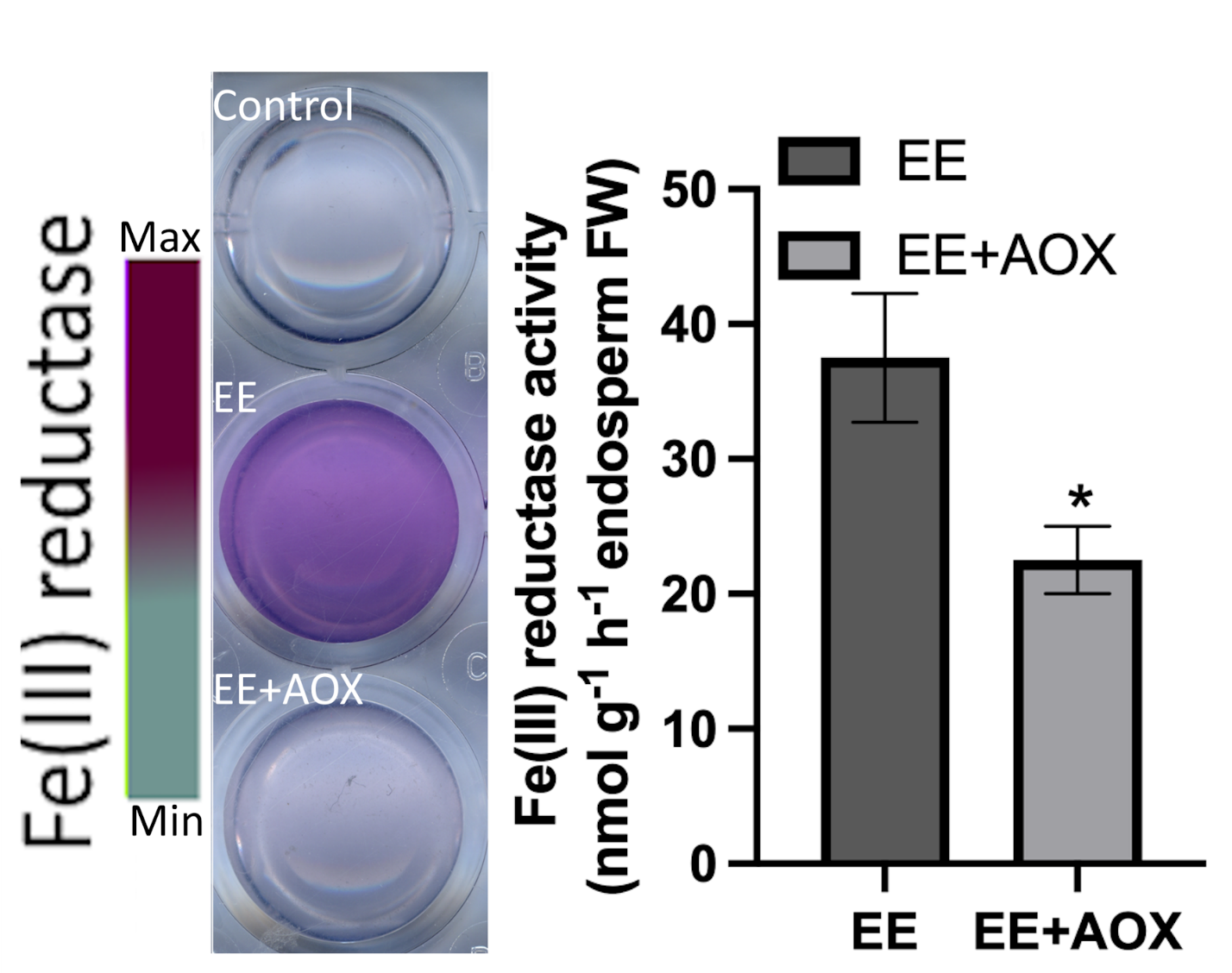

**Figure S6: Imbibed endosperm secretes ascorbate.** Tomato seeds imbibed for one day were cut, and endosperms were isolated and placed in distilled water for 24 hours to collect exudates (EE). The exudates were then mixed with a ferric reductase activity assay solution without (EE) or with ascorbate peroxidase (EE+AOX). The purple color indicates ferric chelate reductase activity. Error bars represent SEM. The asterisk indicates that the corresponding mean of the treatment is significantly different from the mean of the control according to Student’s *t*-test (*P*$<$0.05). The experiment was repeated at least three times with comparable results. AOX: Ascorbate oxidase

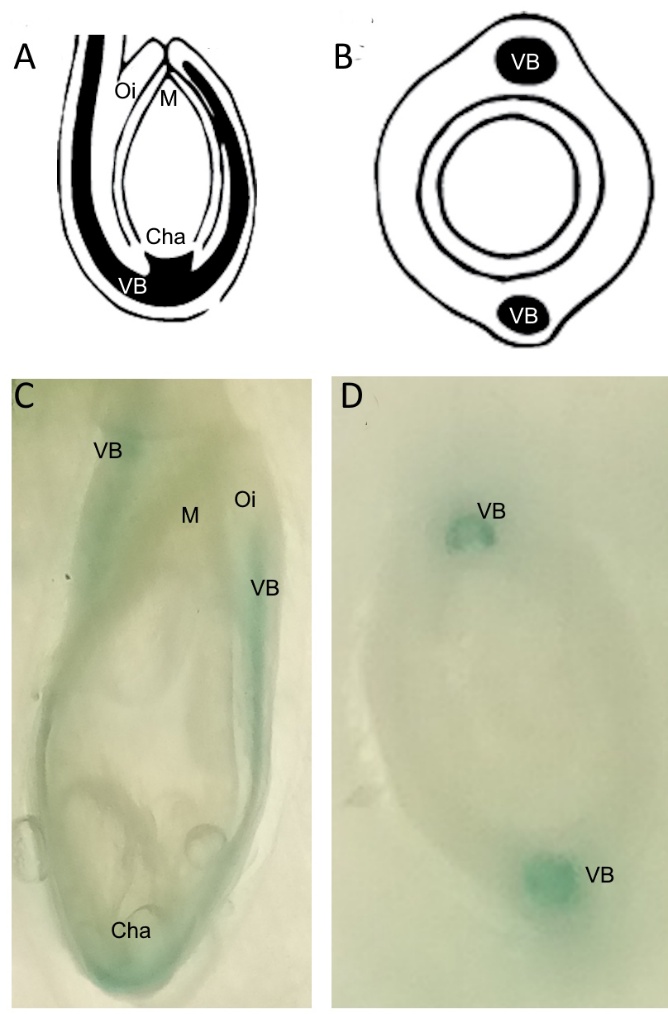

**Figure S7: Variations in chalazal Fe-enriched regions follow variations in vascular invaginations**. A, B; sketches of vascular supply from the mother plant to the developing seed of the Cucurbitaceae family from the side (B) and the top (B), modified from Johri (2013). The black color illustrates vascular bundles that reach the chalaza of the seed and continue inside the outer integument of the sed coat typical of the Cucurbitaceae family. C, D; Perls-stained squash (*Cucurbita pepo*) pieces from the side (C) and from the top (D). Blue coloration shows Fe accumulation. VB: vascular bundle, M: micropylar region, Cha: chalazal region, C: cotyledon, Oi: outer integument of the seed coat. Bar: 50 µm

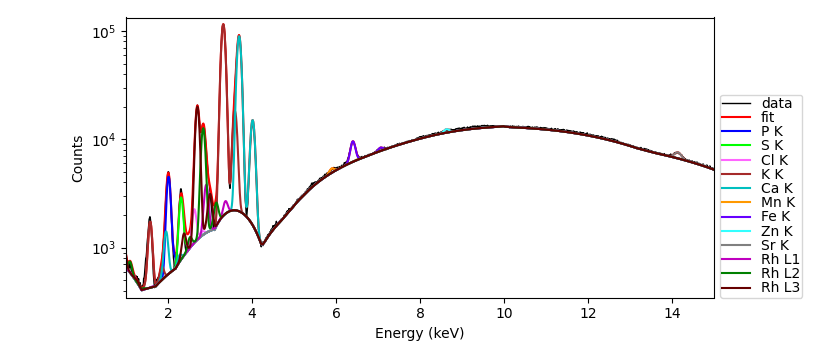

**Figure S8:** Spectrum recorded on citrus fruit. Fit was performed in PyMCA taking into account polychromatic excitation with Rh tube at 30 kV. Spectrum between 100 and 1500 channels was fit, thus scattering peaks are not visible.

**Table S1:** The name of all fruits used in this study

| No | Fruit |
| --- | --- |
| 1 | Strawberry |
| 2 | Parsimmon |
| 3 | Kiwi |
| 4 | Avacado |
| 5 | Pumpkin |
| 6 | Plantain |
| 7 | Papaya |
| 8 | Mango |
| 9 | Pear |
| 10 | Quince |
| 11 | Pepper |
| 12 | Eggplant |
| 13 | Banana |
| 14 | Kumquat |
| 15 | Apple |
| 16 | Tomato |
| 17 | Black mulberry |
| 18 | Lemon |
| 19 | Bean |
| 20 | Orange |
| 21 | Pepino |
| 22 | Grapefruit |
| 23 | Mandarin |
| 24 | Okra |
| 25 | Water melon |
| 26 | Cucumber |
| 27 | Apricot |
| 28 | Citrus |
